# NKG2D signaling regulates IL-17A-producing γδT cells to promote cancer progression

**DOI:** 10.1101/2021.08.26.457761

**Authors:** Sophie Curio, Sarah C. Edwards, Toshiyasu Suzuki, Jenny McGovern, Chiara Triulzi, Nagisa Yoshida, Gustav Jonsson, Teresa Glauner, Damiano Rami, Rachel Violet Purcell, Seth B. Coffelt, Nadia Guerra

## Abstract

γδT cells are unconventional T cells particularly abundant in mucosal tissues that play an important role in tissue surveillance and homeostasis. γδT cell activation is mediated by the T cell receptor composed of γ and δ chains, as well as activating receptors for stress-induced ligands, such as NKG2D. Contrary to the well-established anti-tumor function of γδT cells, recent studies have shown that γδT cells can promote tumor development in certain contexts. However, the mechanisms leading to this disease-promoting role remain poorly understood. Here, we show that mice lacking γδT cells survive longer in a mouse model of intestinal cancer, further supporting their pro-tumoral role. In a surprising conceptual twist, we found that these pro-tumor γδT cells are regulated by NKG2D signaling, a receptor normally associated with cancer cell killing. Germline deletion of *Klrk1*, the gene encoding NKG2D, reduced the frequency of γδT cells in the tumor microenvironment and delayed tumor progression. We further show that blocking NKG2D reduces the capability of γδT cells to produce IL-17A in the pre-metastatic lung and that co-culture of lung T cells with NKG2D ligand-expressing tumor cells specifically increases the frequency of γδT cells. Together, these data support the hypothesis that in a tumor microenvironment where NKG2D ligands are constitutively expressed, γδT cells accumulate in an NKG2D-dependent manner and drive tumor progression by secreting pro-inflammatory cytokines, such as IL-17A.

## Introduction

Cancers affecting mucosal tissues, such as the intestine or the lung, present a global health burden, together making up more than 20% of deaths caused by cancer annually (Sung et al., 2021). Mucosal tissues contain large numbers of unconventional lymphocytes, including γδT cells, which are involved in tissue surveillance and play an important role in the early response against infection and cancer (Hayday, 2009). Indeed, skin-resident γδT cells, activated by stress ligands through the immunoreceptor NKG2D, protect against chemically induced carcinogenesis (Girardi et al., 2001; Strid et al., 2008). Nonetheless, other γδT cell subsets can contribute to malignancies in certain contexts. γδT cells in the lung play a critical role in supporting pulmonary metastasis of p53-deficient mammary tumors, and these lung-resident cells are activated in the pre-metastatic niche by tumors before the spreading of cancer cells in the lung (Coffelt et al., 2015; Wellenstein et al., 2019). In addition to the observation that γδT cells accumulate in several human tumors (Meraviglia et al., 2017; Patil et al., 2016; Rong et al., 2016; Wu et al., 2014), they also contribute to intestinal tumorigenesis in mice (Housseau et al., 2016; Marsh et al., 2012) and promote primary lung cancer in a microbiota-dependent manner (Jin et al., 2019). Yet, the molecular pathways and tissue environments leading to this pro-tumor role of γδT cells remain poorly understood.

NKG2D is a potent immunoreceptor expressed on various innate and adaptive immune cells including γδT cells (Lanier, 2015; Spits et al., 2016). Ligands for NKG2D are stress-induced molecules typically absent from healthy tissue (Diefenbach et al., 2001). Upon binding to its ligands, NKG2D elicits a strong immune response, resulting in the secretion of cytokines and cytotoxic molecules to induce target cell apoptosis. The presence of NKG2D ligands on a large variety of tumors identified the NKG2D/NKG2D ligand axis as an important player in anti-tumor immunity. This has been demonstrated in various *in vitro* killing assays and mouse models of cancer (Cerwenka et al., 2001; Diefenbach et al., 2001; Guerra et al., 2008). This makes NKG2D a promising candidate for use in immunotherapy with several clinical trials exploring NKG2D as a potential chimeric antigen receptor (CAR), to specifically eliminate NKG2D ligand-expressing tumor cells (Baumeister et al., 2018; Xiao et al., 2019). In addition to its anti-tumor effect, NKG2D can contribute to various inflammatory disorders, including inflammatory bowel diseases (Allez et al., 2007; Meresse et al., 2004) and allergic asthma (Farhadi et al., 2014). In the context of inflammation-driven cancer, NKG2D-expressing cells can ultimately lead to cancer progression through their ability to sustain tissue inflammation and cause tissue damage. This was evidenced in a model of chemically induced hepatocellular carcinoma where NKG2D ligands are highly expressed in both tumor and non-tumor adjacent liver tissues (Sheppard et al., 2017, 2018). Whether this pro-tumor effect is limited to the liver tissue or associated with other cancer types driven by inflammation remains to be demonstrated.

In this study, we further dissect the tissue environments and mechanisms leading to the protumor function of γδT cells and NKG2D and show that intestinal tumor progression is delayed in both γδT cell- and NKG2D-deficient mice. NKG2D-deficiency was associated with a marked reduction of γδT cells in the tumor microenvironment (TME). Further, NKG2D-expressing γδT cells displayed an increased ability to produce IL-17A in diseased mice. Importantly, blocking NKG2D reduced IL-17A production in the pre-metastatic lung of a mammary tumor model, extending our observation to an additional cancer site. Lastly, we demonstrate that lung γδT cells expand in an NKG2D-dependent manner *in vitro*. Together, we address the controversy of NKG2D-mediated tumor progression and suggest a pro-tumor function for NKG2D^+^ γδT cells in mucosal tissues.

## Results & Discussion

### γδT cells and NKG2D promote intestinal cancer progression

To investigate the function of γδT cells in intestinal cancer and test the hypothesis that these cells contribute to tumor progression in an NKG2D-dependent manner by creating an inflammatory, tumor-promoting environment, we employed a cancer model driven by the complete loss of the *Apc* gene specifically in intestinal epithelial cells: the *Villin-Cre^ERT2^;Apc^F/+^* model. Similar to patients with mutations in the *APC* gene, these mice are predisposed to intestinal adenoma formation. In this model, γδT cells are the mains producers IL-17A - a cytokine that promotes disease progression in human patients (Grivennikov et al., 2012) - rather than CD4^+^ or CD8^+^ T cells (Supplementary Figure 1a). We crossed *Villin-Cre^ERT2^;Apc^F/+^* mice with mice carrying a germline deletion of the *Tcrd* gene, resulting in the complete loss of γδT cells (*Villin-Cre^ERT2^;Apc^F/+^;Tcrd*^−/−^). Tumor-bearing mice lacking γδT cells survived longer than control mice (Figure 1a), confirming a pro-tumorigenic role for IL-17A-producing γδT cells in *Apc*-deficient intestinal tumors.

**Figure 1.**
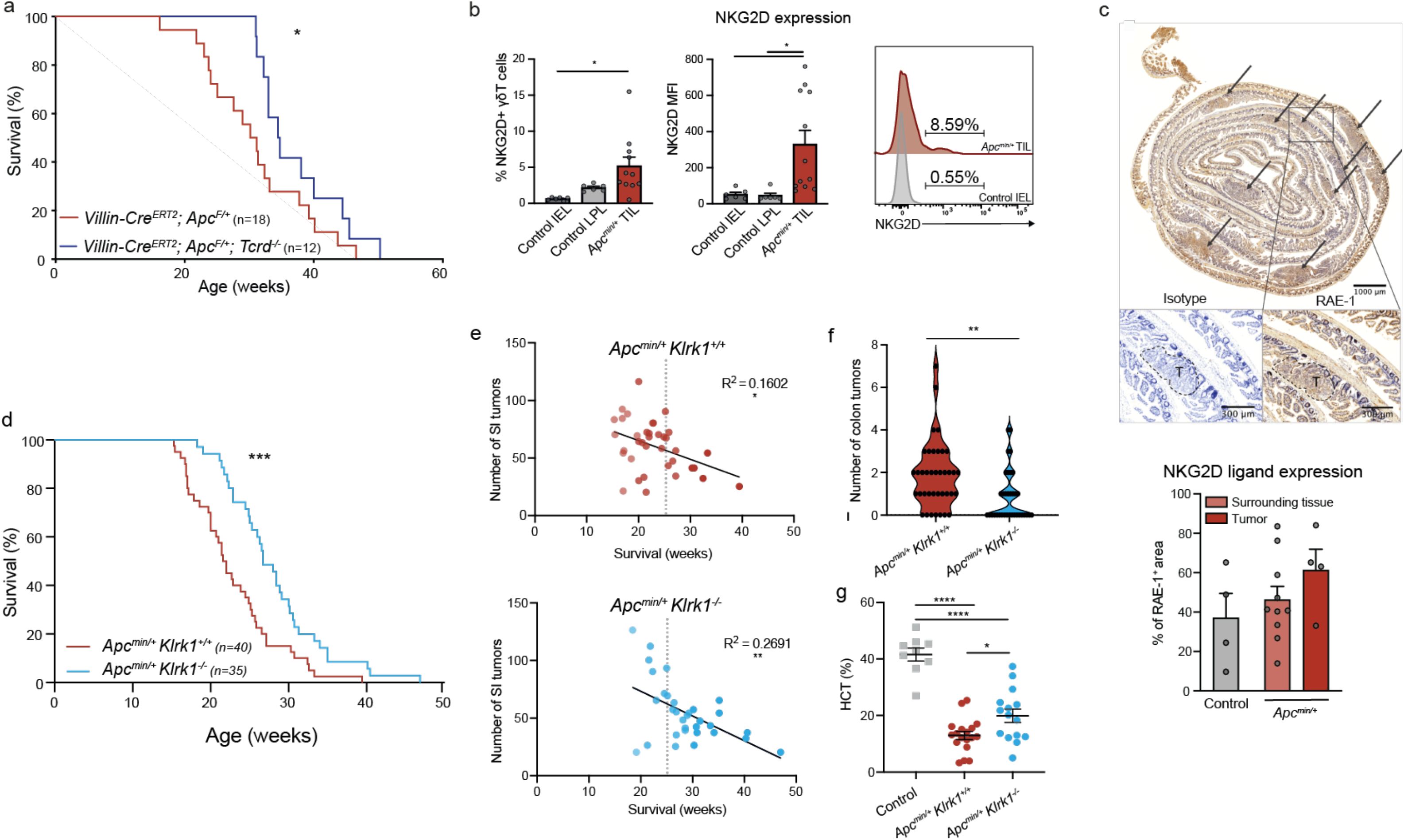
Accelerated tumorigenesis in the absence of γδT cells and NKG2D. | a. Survival of *Villin-Cre^ERT2^;Apc^F/+^* compared to γδT cell-deficient *Villin-Cre^ERT2^; Apc^F/+^;Tcrd*^−/−^ mice. b. Frequencies (left), MFI (middle) and representative histograms (right) of NKG2D-expressing γδT cells in the healthy SI IEL and LPL (n = 6) from naïve mice compared to TIL isolated from SI tumors of *Apc^min/+^* mice (n = 8-15) at 18-20 weeks. c. Representative image of RAE-1 staining in the small intestine of *Apc^min/+^* mice including the unstained isotype control (top) and quantification of RAE-1^+^ area in wildtype control mice (n = 4) compared to tissue surrounding the tumor (n = 10) and the tumor (n = 4) of *Apc^min/+^* mice determined by immunohistochemistry. T = tumor. d. Kaplan-Meier survival analysis of *Apc^min/+^;Klrk1*^+/+^ (n = 40) and *Apc^min/+^;Klrk1*^−/−^ (n = 35) mice. e. Correlation between number of small intestinal (SI) tumors and survival in *Apc^min/+^;Klrk1*^+/+^ (top, red) and *Apc^min/+^;Klrk1*^−/−^ (bottom, blue) mice at disease endpoint. Data points represent individual mice. f. Number of tumors in the colon of *Apc^min/+^;Klrk1*^+/+^ (n = 40) and *Apc^min/+^;Klrk1*^−/−^ mice (n = 35) at disease endpoint. g. Hematocrit (HCT) measures from the blood of healthy *Klrk1*^+/+^ control mice (n = 8), *Apc^min/+^;Klrk1*^+/+^ (n = 18) and *Apc^min/+^;Klrk1*^−/−^ (n = 15) mice at disease endpoint. Data points represent individual mice. SI = small intestine, LPL = lamina propria lymphocytes, IEL = intraepithelial lymphocytes, HCT = hematocrit. Bars represent mean ± SEM. Significance was determined using Log-rank (Mantel-Cox) test (a, c), Mann-Whitney U or unpaired t-test following Shapiro-Wilk normality test (b, c, f and g) and linear regression (d). * p ≤ 0.05, ** p ≤ 0.01, *** p ≤ 0.001 and **** p ≤ 0.0001.

Previous studies investigating the function of γδT cells in intestinal tumorigenesis have shown a disease-promoting role for γδT cells in *Apc^min/+^* mice (Housseau et al., 2016; Marsh et al., 2012), which harbor a point mutation in the *Apc* gene and present with a similar phenotype as the *Villin-Cre^ERT2^;Apc^F/+^* mice. In *Apc^min/+^* mice, the number of NKG2D^+^ γδT cells increases in the intestinal epithelium (Marsh et al., 2012). To corroborate these findings, we determined frequencies of NKG2D^+^ γδT cells within the tumor microenvironment of our own *Apc^min/+^* cohort and compared these frequencies to healthy tissue. In line with previous studies, we found that NKG2D^+^ γδT cells were increased more than eight-fold among tumor-infiltrating lymphocytes (TIL) compared to intestinal epithelial cells (IEL) and more than two-fold when comparing to lamina propria lymphocytes (LPL) of healthy control mice (Figure 1b). In addition to a change in frequencies, we found an increase in mean fluorescence intensity (MFI) of NKG2D on γδT cells present in the tumor microenvironment of *Apc^min/+^* mice compared to healthy tissue (Figure 1b middle and right). We questioned whether NKG2D ligands, which are constitutively expressed on healthy human intestinal epithelium (Groh et al., 1996), are present on tumors of *Apc^min/+^* mice by measuring the expression of the mouse RAE-1. We confirmed that RAE-1 is expressed in the intestine of age-matched healthy control mice and show that RAE-1 is present at significant levels in tumors as well as tumor-surrounding tissue of *Apc^min/+^* mice (Figure 1c). Taken together, the increased frequency of NKG2D^+^ γδT cells within the tumor microenvironment as well as the presence of NKG2D ligand suggests that NKG2D^+^ γδT cells contribute to the immune response in *Apc^min/+^* mice.

To study the function of NKG2D (encoded by *Klrk1*) in the development of intestinal tumors, *Apc^min/+^* mice were intercrossed with *Klrk1*^+/−^ mice to generate NKG2D-sufficient (*Apc^min/+^;Klrk1*^+/+^) and NKG2D-deficient (*Apc^min/+^;Klrk1*^−/−^) mice that were assessed for survival and tumor burden. *Apc^min/+^;Klrk1*^−/−^ mice displayed significantly increased survival compared to *Apc^min/+^;Klrk1*^+/+^ littermates (Figure 1d), a delay in development of small intestinal tumors (Figure 1e), as well as a twofold decrease in colonic tumors at disease endpoint (Figure 1f). Hematocrit (HCT) was measured as an additional read-out of disease severity. Anemia is a direct consequence of the high numbers of multiple polyps in the small intestine (SI), which can exceed 100 tumors per mouse, leading to significant blood loss. Blood collected at disease endpoint showed lower HCT levels in *Apc^min/+^;Klrk1*^+/+^ mice compared to *Apc^min/+^;Klrk1*^−/−^ mice (Figure 1g), further supporting the observations that tumor progression is enhanced in the presence of NKG2D. Collectively, these data show that γδT cells infiltrating intestinal tumors express a significant amount of IL-17A and NKG2D and these cells drive disease progression in *Apc*-mutated models of intestinal tumorigenesis.

### NKG2D regulates pro-tumorigenic γδT cells in intestinal tumors

To assess how NKG2D signaling influences immune cell infiltration and activation in intestinal tumors, we compared the frequencies of tumor-infiltrating immune cell subsets in *Apc^min/+^;Klrk1*^+/+^ and *Apc^min/+^;Klrk1*^−/−^ mice. We found that the frequency of T cells was significantly higher in tumors from *Apc^min/+^;Klrk1*^+/+^ mice compared with tumors from *Apc^min/+^;Klrk1*^−/−^ mice, mainly due to an increased proportion CD8^+^ T cells and γδT cells in NKG2D-sufficient mice (Figure 2a,b). These data suggest that NKG2D signaling influences infiltration and/or accumulation of these specific T cell subsets.

**Figure 2.**
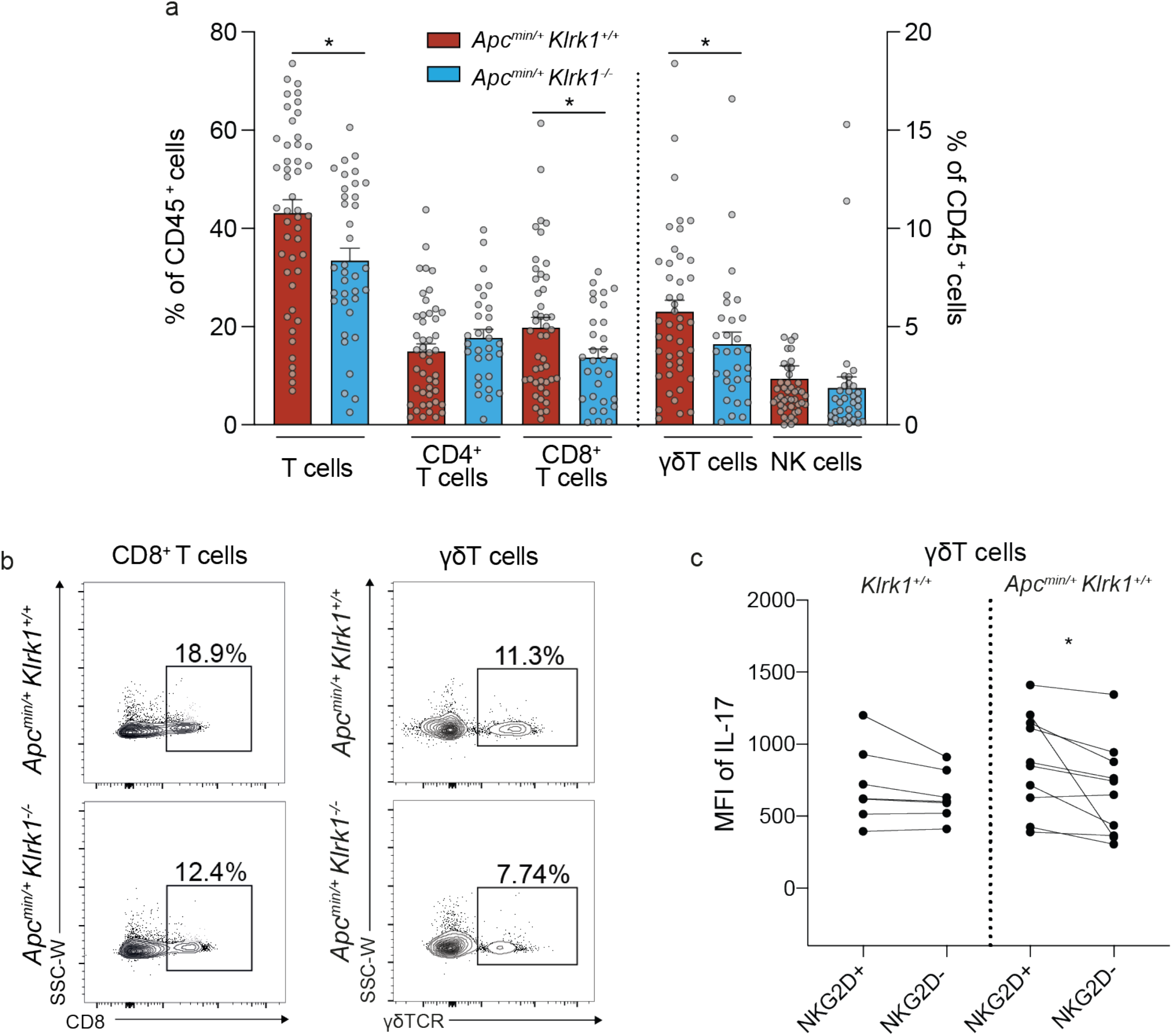
NKG2D regulates immune cell infiltration and cytokine production in the tumor microenvironment. | a. Average frequencies of T cells (CD3^+^), CD4^+^ T cells (CD3^+^CD4^+^), CD8^+^ T cells (CD3^+^CD8^+^), γδT cells (CD3^+^γδTCR^+^) and NK cells (CD3^-^NK1.1^+^) in the TME of SI tumors of *Apc^min/+^;Klrk1*^+/+^ (n = 41-47) compared to *Apc^min/+^;Klrk1*^−/−^ mice (n = 29-33) at disease endpoint. b. Representative flow cytometry plots depicting CD8^+^ and γδT cells in the in the TME of SI tumors of *Apc^min/+^;Klrk1*^+/+^ compared to *Apc^min/+^;Klrk1*^−/−^ mice. c. Direct comparison of IL-17A MFI in NKG2D^+^ and NKG2D^−^ γδT cells, gated on live CD3^+^ lymphocytes, isolated from the MLN of *Klrk1*^+/+^ (n = 6) and *Apc^min/+^;Klrk1*^+/+^ mice at 18-20 weeks (n = 10). TME = tumor microenvironment, SI = small intestine, MLN = mesenteric lymph nodes. Significance was determined using Mann-Whitney U or unpaired t-test following Shapiro-Wilk normality test (a and c) or paired t-test (d). Bars represent mean ± SEM. * p ≤ 0.05. Data points represent individual mice.

IL-17A, a pro-inflammatory cytokine known to drive intestinal tumorigenesis in the *Apc^min/+^* mouse model (Chae et al., 2010), is produced by γδT cells. Thus, we hypothesized that NKG2D regulates IL-17A expression, thereby contributing to the inflammatory milieu and tumor progression. We measured the quantity of IL-17A produced by NKG2D^+^ γδT cells (mean fluorescence intensity, MFI) localized in the MLN of tumor bearing and healthy mice and found that was it was higher than the MFI measured in NKG2D^−^ γδT cells (Figure 2c). These data support the concept that pro-tumoral IL-17A-producing γδT cells are regulated in part by NKG2D signaling.

### NKG2D signaling stimulates pro-tumorigenic IL-17A-producing γδT cells

To further probe the role of NKG2D on IL-17^+^ γδT cells, we used a mammary tumor model where IL-17A-producing γδT cells drive lung metastasis (Coffelt et al., 2015; Wellenstein et al., 2019). IL-17A-producing γδT cells in the lung play a critical role in supporting pulmonary metastasis of p53-deficient mammary tumors, and these lung-resident cells are activated in the pre-metastatic niche by tumors before the arrival of cancer cells in the lung (Coffelt et al., 2015; Wellenstein et al., 2019). Mice bearing *K14-Cre;Brca1^F/F^;Trp53^F/F^* (KB1P) mammary tumor transplants (Liu et al., 2007) develop lung metastases and present a good model for studying immune remodeling of the pre-metastatic lung (Millar et al., 2020). Similar to the intestine, NKG2D is highly expressed on immune cells in the lung, in particular on lung γδT cells and NK cells (Figure 3a,b). We therefore questioned whether NKG2D signaling could impact IL-17A expression by lung γδT cells. KB1P tumor-bearing mice were treated with IgG control antibodies or anti-NKG2D blocking antibodies and cytokine expression by γδT cells and other lymphocytes in the pre-metastatic lung (i.e. lung tissue conditioned by primary mammary tumors) was measured. Blocking NKG2D resulted in reduced IL-17A expression in γδT cells when compared with controls (Figure 3c,d), indicating that NKG2D signaling positively regulates pro-metastatic IL-17A-producing γδT cells in the lung.

**Figure 3.**
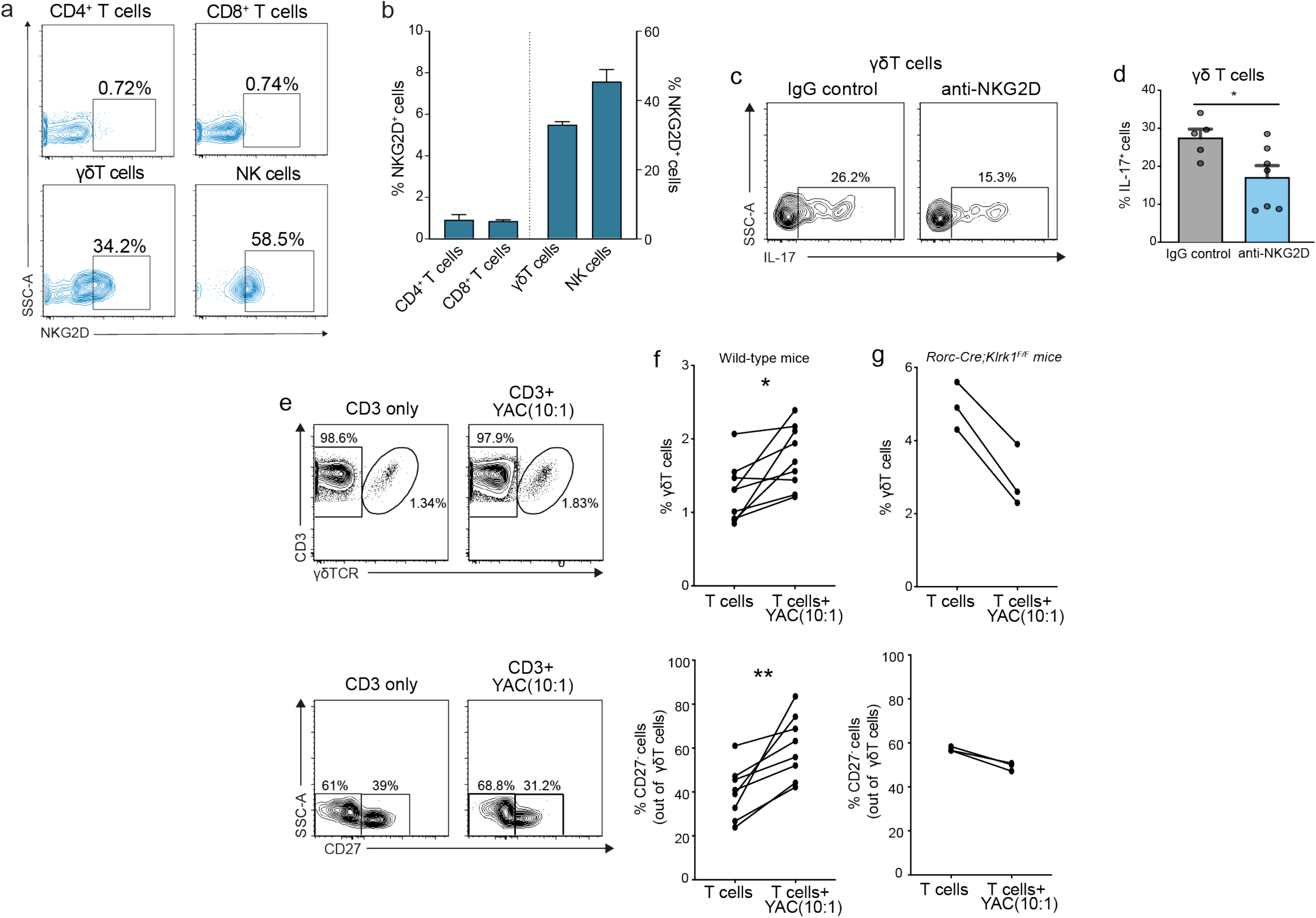
NKG2D signaling regulates IL-17A-producing T cells. | a. Representative flow cytometry dot plots depicting NKG2D-expressing cells in the lung. b. Average frequencies of NKG2D-expressing cells among T cells and NK cells in the lung (n = 3). c. Representative flow cytometry dot plots depicting IL-17A staining on γδT cells in mice bearing KB1P tumor transplants treated with isotype control IgG antibody or anti-NKG2D antibody. d. Average frequencies of IL-17A-producing γδT cells, gated on live CD3^+^ T cells in mice bearing KB1P tumor transplants treated with the isotype control IgG (n = 5) antibody or anti-NKG2D (n = 7) antibody. e. Representative flow cytometry dot plots of γδTCR (top, gated on CD3^+^ live lymphocytes) and CD27 expression (bottom, gated on γδTCR^+^CD3^+^ live lymphocytes) before and after co-culture of T cells isolated from the lung with NKG2D ligand-expressing YAC-1 cells. f. Average frequencies of γδT cells (top) and CD27^−^ γδT cells (bottom) isolated from wild-type mice before and after co-culture of T cells isolated from the lung with YAC cells (n = 8). Each line represents one individual mouse. g. Frequencies of *Rorc-Cre;Klrk1^F/F^* γδT cells (top) and CD27^−^ γδT cells (bottom) before and after co-culture of T cells isolated from the lung with YAC cells (n = 3). Each line represents one individual mouse. Bars represent mean ± SEM. Significance was determined using Mann-Whitney U or unpaired t-test following Shapiro-Wilk normality test (b) or paired t-test (d). * p ≤ 0.05, ** p ≤ 0.01.

To gain further insight into the regulation of γδT cells by NKG2D, we performed co-culture experiments where CD3^+^ T cells from the lungs of naïve mice were incubated with the NKG2D ligand (RAE-1)-expressing lymphoma cell line, YAC-1. In this assay, the proportion of γδT cells (Figure 3e,f), but not other T cells (data not shown) increased after interaction with YAC-1 cells. This was specifically due to an increase in CD27^−^ γδT cells, which are capable of producing IL-17A (Figure 3e,f). To show that this increase in CD27^−^ γδT cells by RAE-1-expressing YAC-1 cells was directly dependent on NKG2D signaling, we generated mice whose IL-17A-producing γδT cells lack NKG2D expression. We crossed *Rorc-Cre* mice with *Klrk1^F/F^* mice so that activation of RORγt, the master transcription factor of the *Il17a* gene, deletes *Klrk1* by Cre recombinase. Lung CD3^+^ T cells from *Rorc-Cre;Klrk1^F/F^* mice were then co-cultured with RAE-1-expressing YAC-1 cells. However, in contrast with NKG2D-proficient γδT cells (Figure 3f), the proportions of γδT cells and NKG2D-deficient *Rorc-Cre;Klrk1^F/F^* CD27^−^ γδT cells failed to increase after co-culture with YAC-1 cells (Figure 3g). These data demonstrate that lung NKG2D^+^CD27^−^ γδT cells selectively expand when cultured with RAE-1-expressing YAC-1 cells. Collectively, these data indicate that NKG2D signaling enhances activation of γδT cells, which are a source of pro-inflammatory IL-17A and contribute to tumor progression.

In this study, we show that both γδT cells and NKG2D contribute to the development and progression of mucosal tumors. Germline deletion of *Tcrd* as well as *Klrk1* result in an improved survival in a model of intestinal cancer. We demonstrate a reduced frequency of tumor-infiltrating γδT cells in tumors of NKG2D-deficient mice and show that co-culture of T cells isolated from the lung with NKG2D ligand-expressing tumor cells specifically increases the frequency of γδT cells. Lastly, we confirm the relationship between NKG2D and IL-17A by showing that antibody-mediated blocking of NKG2D specifically reduces the frequency of IL-17A-producing γδT cells in the lung.

Our findings reveal a new molecular pathway of γδT cells-mediated tumor progression: while γδT cells have the potential to recognize tumor cells and initiate an anti-tumor response through the engagement of NKG2D, we provide further evidence for a pro-tumorigenic role in inflammation-driven mucosa-associated cancers. These findings underline the need to better understand the function of tumor-infiltrating γδT cells and NKG2D-expressing immune cells, in particular in light of recent clinical studies using γδT cell- and NKG2D-based approaches in immunotherapy (Lonez et al., 2018; Lu et al., 2015; Murad et al., 2018). While engagement of NKG2D can potentially lead to tumor regression by unleashing an anti-tumor immune response, we show that expression of NKG2D and its ligands in mucosal cancers can activate tumor-promoting immune cells, such as IL-17A-producing γδT cells. We have previously shown that NKG2D-expressing CD8^+^ T cells promote cancer in a model of liver cancer (Sheppard et al., 2017). Here, we extend these findings of a pro-tumor role for NKG2D to intestinal cancer and lung metastasis, and show that disease progression is mediated through distinct mechanisms. Current studies using anti-NKG2D antibodies to treat Crohn’s Disease patients (Vadstrup and Bendtsen, 2017), as well as ongoing trials on NKG2D CAR-T cells in colorectal cancer patients (Celyad, NCT03310008) will provide crucial information on the function of NKG2D in the context of intestinal inflammation and cancer. Together, we unravel a novel role of NKG2D-expressing γδT cells in mucosal immunity and tumorigenesis and uncover a paradoxical role of these cells in cancer. Our data highlights the need to better understand the function of NKG2D^+^ γδT cells and NKG2D ligand expression in the tumor microenvironment, including tissue-specific immune responses.

## Methods

### Mice

NKG2D-heterozygous (*Klrk1^+/-^*) (Guerra et al., 2008) mice were crossed with *Apc^min/+^* mice to generate *Klrk1^+/+^;Apc^min/+^*, *Klrk^−/−^;Apc^min/+^*, *Klrk1^+/+^;Apc*^+/+^ and *Klrk1^−/−^;Apc^+/+^* mice and to study the impact of NKG2D-deficiency in the intestine. Studies on the lung were performed on *Klrk1*^+/−^ mice provided by Dr. Bojan Polic (University of Rijeka School of Medicine, Croatia). *Villin-Cre^ERT2^;Apc^F/+^;Tcrd^−/−^* mice were generated by crossing *Villin-Cre^ERT2^;Apc^F/+^* mice with *Tcrd*^−/−^ mice. The alleles used were as follows: *Villin-Cre^ERT2^* ^42^, *Apc^F^* (Marjou et al., 2004; Shibata et al., 1997) and *Tcrd*^−/−^ (Itohara et al., 1993). *Villin-Cre^ERT2^* experiments were performed on a mixed background (N8 for C57BL/6J). Recombination by *Villin-Cre^ERT2^* was induced with one intraperitoneal (i.p.) injection of 80 mg/kg tamoxifen when mice reached 20 g. The health status of *Apc^min/+^* and *Villin-Cre^ERT2^;Apc^F/+^* mice was checked and evaluated frequently. Disease severity was assessed using a scoring scheme that included parameters such as appearance, natural behavior, provoked behavior, body condition, and tumor score. For hematocrit (HCT) measurement, blood was collected from the tail vein of live mice into Eppendorf tubes containing 100 μl 0.5 M EDTA to prevent clotting. Blood parameters, including hematocrit. were measured using the XN-350 (Sysmex). Mice were humanely euthanized when they had reached the experimental endpoint.

*Klrk1^F/F^* mice were provided by Dr. Bojan Polic (University of Rijeka School of Medicine, Croatia) and crossed with *Rorc-Cre* mice (JAX) to generate *Rorc-Cre;Klrk1^F/F^* mice.

*K14-Cre;Brca1^F/F^;Trp53^F/F^* (KB1P) mice were a gift from Dr. Jos Jonkers (Netherlands Cancer Institute). KB1P mice were maintained on FVB background and generation of this strain has previously been described, where a human K14 gene promoter drives *Cre* recombinase transgene expression, resulting in LoxP *Trp53* and *Brca1* specific deletion in the mammary epithelium(Liu et al., 2007). This genetically engineered mouse model resembles human triple negative breast cancer and mice develop single mammary tumors at approximately 25-30 weeks of age. *Cre* recombinase negative (*Cre^-^*) female littermates were used as wild-type (WT) controls. Female FVB mice (8-10 weeks of age) were obtained from Charles River. Mice were palpated twice per week from 12 weeks of age for mammary tumors and perpendicular tumor diameters were measured with a caliper. Mice were sacrificed once a tumor reached >1 cm in any direction.

Mice were bred and maintained in the animal facility at Imperial College London or the Cancer Research Beatson Institute (Glasgow, UK) in a specific pathogen-free environment. Work was carried out in compliance with the British Home Office Animals Scientific Procedures Act 1986 and the EU Directive 2010 and sanctioned by Local Ethical Review Process (PPL 70/8606 and 70/8645).

### Blockade of NKG2D in vivo

Female FVB/n mice at 8-10 weeks old (purchased from Charles River) were orthotopically transplanted in their mammary fat pat with 1×1 mm tumor pieces from syngeneic *K14-Cre;Brca1^F/F^;Trp53^F/F^* (KB1P) mice. Once tumors reached 1cm, animals were injected intraperitoneally with a single dose of 200 mg anti-NKG2D (Clone HMG2D, BioXCell) on day 1 followed by individual injections of 100 mg on two consecutive days. Control mice followed the same dosage regime with Armenian hamster IgG isotype control (Clone Polyclonal, Bio X Cell). Tumor growth was measured by calipers and mice were euthanized one day after final antibody injection.

### Tissue processing

Tumors isolated from KB1P mice were collected in PBS, cut into small pieces, and resuspended in Dulbecco’s Modified Eagle Medium F12 (DMEM) containing 30% fetal calf serum (FCS) and 10% dimethyl sulfoxide and stored at −150 °C. Lungs, lymph nodes (LN) (axillary, brachial, mesenteric), and/or the spleen were isolated in ice-cold phosphate-buffered saline (PBS). Lungs were mechanically dissociated by using a scalpel and transferred to collagenase solution consisting of DMEM supplemented with collagenase D (Roche, 1 mg/ml) and DNase1 (ThermoFisher, 25 μg/ml). Enzymatic dissociation was assisted by heat and mechanical tissue dissociation, using the gentleMACS Octo Dissociator (Miltenyi Biotec) (run name: 37C_m_LDK), according to the manufacturer’s dissociation protocol. The lung cell suspension was filtered through a 70 μm cell strainer using a syringe plunger and enzyme activity was stopped by addition of 2 ml of FCS followed by 5 ml DMEM medium supplemented with 10% FCS, L-glutamine (2 mM, ThermoFisher) and penicillin (10000 U/ml) / streptomycin (10000 μg/ml, ThermoFisher). Spleen and LN were processed and filtered through a 70 μm cell strainer using a syringe plunger, and the tissue was flushed through with PBS containing 0.5% Bovine Serum Albumin (BSA). Cell suspension were centrifuged at 201 x g for 5 min, and supernatant was discarded. Cells were resuspended in PBS containing 0.5% BSA and cell number was determined using a hemocytometer.

The small intestine (SI) and tumor from control mice (age-matched *Villin-Cre^ERT2^* negative;*Apc^F/+^* mice) and *Villin-Cre^ERT2^;Apc^F/+^* mice were collected in PBS, cut into small pieces using a McIlwain tissue chopper (Campden Instruments Ltd), and digested by heat and mechanical tissue dissociation using the gentleMACS Octo Dissociator (Miltenyi Biotec) (run name: 37C_m_TDK_1), according to the manufacturer’s dissociation protocol. Cells were resuspended in PBS containing 0.5% BSA.

Small intestine and colon were harvested from healthy *Klrk1^+/+^, Klrk1*^−/−^, and *Apc^min/+^* mice. Once the caecum was removed, the small intestine was separated into duodenum, jejunum, and ileum. Data presented in this study relate to tumors isolated from the ileum unless otherwise stated. Remaining fat, as well as the gut content, was removed from the intestine before it was flushed with PBS. For tumor cell isolation, the tumor was carefully excised avoiding cutting through tumors. Mucus was removed, and tumor cells were counted before they were carefully removed from the intestinal tissue and minced using a scalpel. Tissue was transferred to a 1.5 ml tube and 1 ml of RPMI-1640 containing 5% FCS, 25 mM HEPES, 150 U/ml collagenase IV (Sigma Aldrich), and 50 U/ml DNase (Roche) were added. Tubes were shaken in a horizontal position on an incubator shaker (37 °C, 200 rpm, 30 minutes). Dissociated tissue was transferred to a 50 ml tube through a 100 μm cell strainer. To further dissociate the tissue, it was pushed through the filter using the plunger of a 2 ml syringe. To inhibit enzyme activity, 1 ml of PBS containing 10% FCS and 5 mM EDTA was added to the 1.5 ml tube and then transferred to the 50 ml tube. Cells were centrifuged for 10 minutes at 500 x g and resuspended in RPMI-1640 containing 5% FCS.

To isolate intraepithelial lymphocytes, the tissue was cut into small pieces and transferred to a 15 ml tube containing 10 ml of HBSS containing 1 mM DTT, 2% FCS, 100 U/ml penicillin and 100 μg/ml streptomycin. Tubes were vortexed for 30 s and the solution was replaced. Tubes were shaken in a horizontal position on an incubator shaker (37 °C, 200 rpm, 20 minutes), which resulted in the removal of the mucus layer. Using a 100 μm cell strainer, the tissue was then transferred to a new 15 ml tube containing 10 ml of HBSS containing 1 mM EDTA, 2% FCS, 100 U/ml penicillin, and 100 μg/ml streptomycin. It was again shaken in a horizontal position (37 °C, 200 rpm, 15 minutes), before it was filtered through a 100 μm strainer. The liquid was taken up and the tissue transferred back to the 15 ml tube and 10 ml of fresh solution was added. This step was repeated two or three times for large and small intestine, respectively, and resulted in the removal of all epithelial cells. This resulted in a single cell suspension, which was then centrifuged for 10 minutes at 500 x g and resuspended in RPMI-1640 containing 5% FCS.

### Magnetic-activated cell sorting to isolate T cells

To isolate T cells from lung, LN and spleen, single cell suspensions were obtained as previously described. CD3^+^ cell enrichment was achieved by MojoSort negative selection using MojoSort mouse CD3^+^ T-cell Isolation Kit (BioLegend) according to manufacturer’s instructions. In brief, 1×10^7^ cells were resuspended in 100 μL MojoSort buffer and cells were incubated with 10 μL of the biotin-antibody cocktail in a 5 ml polypropylene tube and incubated for 15 min on ice. Ten μL of streptavidin nanobeads were added and mixed and incubated on ice for further 15 min. Two and a half ml of MojoSort buffer was added and the tube placed in the magnet for 5 min to allow magnetically labelled cells to bind to the tube and magnetic separator. The untouched CD3^+^ cells were collected by decanting the liquid into a new tube. Cells after enrichment were counted with a hemocytometer and plated at a density of 1×10^6^ cells/ml in round bottom 96-well plates.

### Cell culture

YAC-1 lymphoma cells were maintained in RPMI-1640 medium supplemented with 10% FCS, 100 U/ml penicillin and streptomycin and 2 mM glutamine and kept in incubators at 37 °C under normoxic conditions. Freshly isolated T cells were added at an effector: target (E:T) ratio of 10:1 to YAC-1 cells. Co-cultures were incubated in normoxia at 20% O_2_, 5% CO_2_, and 37 °C for 48 h in complete IMDM medium (supplemented with 10% FCS, L-glutamine (2 mM) and penicillin (10000 U/ml) and streptomycin (10000 μg/ml) and 50 μM b-mercaptoethanol).

### Flow cytometry

Prior to antibody staining, cells were stimulated with PMA (final concentration 600 ng/ml), ionomycin (final concentration 100 ng/ml) and Brefeldin A (final concentration 10 μg/ml) in RPMI-1640 containing 5% FCS, 100 U/ml penicillin and 100 μg/ml streptomycin to enhance cytokine production. Unspecific binding by the Fc receptor was prevented by incubation of anti-mouse CD16+CD32 (BD) for 20 minutes. Dead cells were stained using the LIVE/DEAD Fixable Aqua Dead Cell Stain kit (ThermoFisher) or the Zombie NIR Fixable Viability Dye (BioLegend) before antibodies were added and incubated for 30 minutes. Before staining of intracellular cytokines and transcription factors, cells were permeabilized using the Transcription Factor Fixation/Permeabilization kit (eBioscience). Antibodies to detect intracellular antigens were added and incubated for 30 minutes before cells were washed and transferred to round-bottom polystyrene 5ml tubes through a filter mesh. Samples were acquired on a BD LSR Fortessa and analyzed using FlowJo software. All antibodies used in this study are listed in Supplementary Table 1.

### Immunohistochemistry

Swiss rolls of intestines were prepared according to published protocols(Bialkowska et al., 2016). Tissues were fixed in 10% formalin in water overnight and then transferred to 70% ethanol where it was kept until paraffin embedding. Paraffin blocks were cut into 4 μm sections using a microtome (Leica), placed in a 40 °C water bath, and affixed onto Superfrost Plus Microscope Slides (Thermo Scientific). Prior to staining, slides were deparaffinized and rehydrated. After heat-induced epitope retrieval (HIER), several blocking steps were performed to prevent non-specific binding of antibodies. Primary antibody (Anti-mouse RAE-1 pan-specific antibody, recognizing RAE-1α, β, δ, γ and ε (R&D Systems, Catalogue Number AF1136) or an appropriate isotype-matched control (polyclonal goat isotype IgG, NEB) was added at a final concentration of 10 μg/ml and incubated overnight. After several washing steps, the secondary antibody (10 μg/ml of biotinylatd rabbit anti-goat, Vectorlabs) was added and incubated for 30 minutes. The antigen was revealed using ABC (Vectorlabs) and DAB reagents (Abcam). Slides were counterstained by immersion in Harris Haematoxylin and dehydrated before coverslips were mounted using HistoMount mounting medium (ThermoFisher). Slides were scanned using an AxioScan.Z1 slide scanner (Zeiss). For image analysis, tumors and surrounding tissue were identified and marked using Fiji software (ImageJ). Percentage of positive cells was determined using a macro by comparing specific stained samples to isotype-matched Ig control samples.

### Statistical analysis

Unless stated otherwise, statistical analysis was performed using Python, Pandas, or GraphPad Prism. Shapiro-Wilk tests were performed to test for normality and significance was determined using Mann-Whitey U or unpaired t tests unless stated otherwise. Data visualization was performed using GraphPad Prism (version 8.4.2).

## Author contributions

S.C. designed and performed experiments, analyzed data and wrote the manuscript. S.C.E and T.S. designed and performed experiments, analyzed data and reviewed the manuscript. J.M., C.T. and N.Y. performed experiments and analyzed data. G.J., T.G. and D.R. contributed to some of the experiments. R.P. supervised the microbiota study. N.G. and S.B.C. designed and supervised the studies, analyzed the data and wrote the manuscript.

## Acknowledgements

The authors declare no competing financial interests. We thank S. Sheppard, P. Guermonprez, J. Helft and G. Gorkiewicz for critical reading of the manuscript, B. Polić for providing reagents and insightful discussion, J. Srivastava and J. Rowley from the Imperial College Flow Facility, L. Lawrence from the Imperial College Research Histology Facility and the Blizard Core Pathology Facility for technical support. We thank the Core Services and Advanced Technologies at the Cancer Research UK Beatson Institute (C596/A17196), with particular thanks to the Biological Services and Flow Cytometry Facilities.

This work was supported by the Wellcome Trust RCDF (RCDF088381/Z/09/Z to N.G.), a Wellcome Trust PhD studentship (203948/Z/16A to S.C.), the Stevenson Fund to S.C., Breast Cancer Now (2018JulPR1101 to S.B.C.), Cancer Research UK Glasgow Centre (A25142 to S.B.C.), Wellcome Trust (208990/Z/17/Z to S.B.C.), Marie Curie European Fellowship (GDCOLCA 800112 to T.S.) and Naito Foundation Grant for Research Abroad (to T.S).

---

**Supplementary Figure 1.**
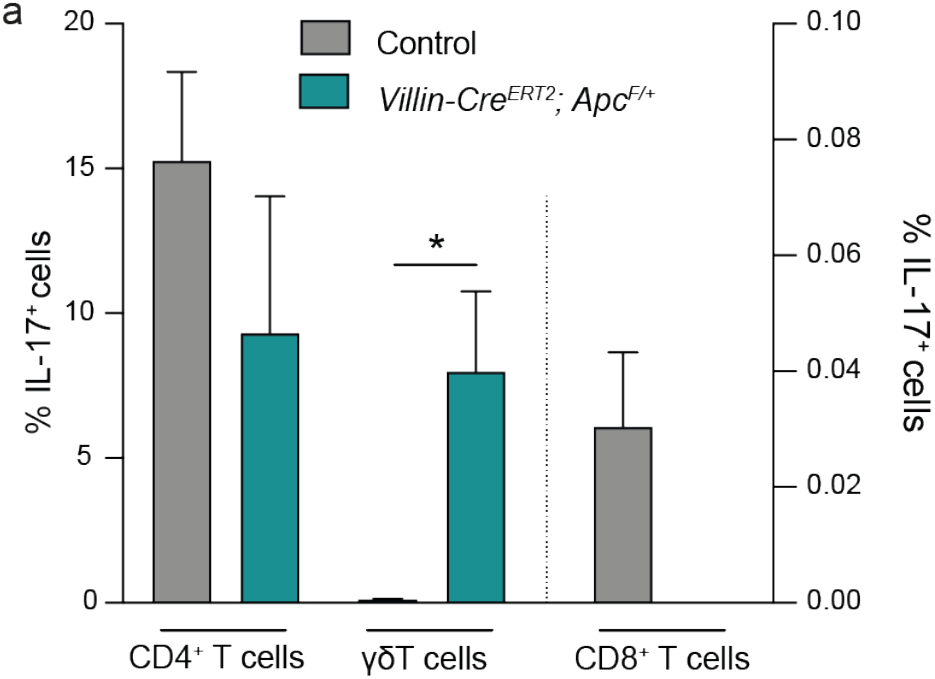
a. Frequencies of IL-17A producing T cells (gated γδTCR^-^CD4^+^, γδTCR^-^ CD8^+^ or CD4^-^CD8^-^γδTCR^+^ following gating on live CD3^+^ lymphocytes) in the *Villin-Cre^ERT2^;Apc^F/+^* mouse model of intestinal cancer at disease endpoint compared to healthy control mice (n = 4). Bars represent mean ± SEM. * p ≤ 0.05.

**Table 1.** Antibodies used in this study.

| Antigen | Conjugate | Clone | Source |
| --- | --- | --- | --- |
| B220 | PE-Cy7 | RA3-6B2 | BioLegend |
| CD11b | APC-eFluor780 | M1/70 | eBioscience |
| CD11b | PE-CF594 | M1/70 | BD |
| CD11b | PerCp-Cy5.5 | M1/70 | eBioscience |
| CD19 | APC-eFluor780 | 1D3 | eBioscience |
| CD19 | BV650 | 6D5 | BioLegend |
| CD27 | BV510 | LG.3A10 | BD |
| CD27 | PE/Dazzle594 | LG.3A10 | BioLegend |
| CD3 | BV650 | 17A2 | BioLegend |
| CD3 | V450 | 17A2 | BD |
| CD3 | PE-CF594 | 145-2C11 | BD |
| CD3 | BUV737 | 145-2C11 | BD |
| CD4 | APC-eFluor780 | GK1.5 | Invitrogen |
| CD4 | BV605 | GK1.5 | BioLegend |
| CD4 | APC-eFluor780 | RM4-5 | eBioscience |
| CD4 | BV711 | RM4-5 | BioLegend |
| CD4 | BUV495 | GK1.5 | BD |
| CD44 | PerCP-Cy5.5 | IM7 | BioLegend |
| CD45 | BV605 | 30-F11 | BioLegend |
| CD45 | AF700 | 30-F11 | BioLegend |
| CD8α | BUV395 | 53-6.7 | BD |
| CD8α | APC-eFluor780 | 53-6.7 | Invitrogen |
| CD8α | PerCp-Cy5.5 | 53-6.7 | BioLegend |
| CD8α | BV785 | 53-6.7 | BioLegend |
| CD8α | BUV395 | 53-6.7 | BD |
| EpCAM | APC-eFluor780 | G8.8 | eBioscience |
| Fixable Aqua Dead Cell Stain Kit | N/A | N/A | ThermoFisher |
| Granzyme B | AlexaFluor-647 | GB11 | BioLegend |
| IFN-γ | PE-Cy7 | XMG1.2 | eBioscience |
| IFN-γ | PE | XMG1.2 | eBioscience |
| IL-17A | FITC | GL3 | eBioscience |
| IL-17A | PE-Cy7 | TC11-18H10.1 | BioLegend |
| IL-17A | PE | eBio17B7 | eBioscience |
| NK1.1 | AlexaFluor-488 | PK136 | BioLegend |
| NK1.1 | PE-Cy7 | PK136 | BioLegend |
| NKG2D | PE/Dazzle594 | CX5 | BioLegend |
| NKG2D | APC | CX5 | BioLegend |
| NKG2D | PE | CX5 | eBioscience |
| NKp46 | BV421 | 29A1.4 | BioLegend |
| PD-1 | BV650 | J43 | BD |
| TCR Vγ1 | PE | 2.11 | BioLegend |
| TCR Vγ4 | APC | UC3-10A6 | BioLegend |
| TCR Vγ4 | PeCy7 | UC3-10A6 | eBioscience |
| TCR Vγ6 | Pacific blue | 1C10-1F7 | Provided by Shinya Hatano and Yasunobu Yoshikai<sup>55</sup> |
| Zombie NIR | N/A | N/A | BioLegend |
| $\gamma\delta$TCR | PerCp-eFluor 710 | GL3 | eBioscience |
| $\gamma\delta$TCR | BV421 | GL3 | BD |
| $\gamma\delta$TCR | APC | GL3 | eBioscience |
| $\gamma\delta$TCR | PE-Cy5 | GL3 | eBioscience |
| $\gamma\delta$TCR | FITC | GL3 | eBioscience |

